# An elasticity-curvature illusion decouples cutaneous and proprioceptive cues in active exploration of soft objects

**DOI:** 10.1101/2020.08.04.237016

**Authors:** Chang Xu, Yuxiang Wang, Gregory J. Gerling

## Abstract

Our sense of touch helps us encounter the richness of our natural world. Across a myriad of contexts and repetitions, we have learned to deploy certain exploratory movements in order to elicit perceptual cues that are optimal and efficient. Such cues help us assess an object’s roughness, or stickiness, or as in this case, its softness. Leveraging empirical experiments combined with computational modeling of skin deformation, we develop a perceptual illusion for softness, or compliance, where small-compliant and large-stiff spheres are indiscriminable. The elasticity-curvature illusion, however, becomes readily discriminable when explored volitionally. This tactile illusion is unique because it naturally decouples proprioceptive cues from those involving identical, cutaneous contact attributes. Furthermore, the illusion sheds light into exactly how we explore soft objects, i.e., by volitionally controlling force, to optimally elicit and integrate proprioceptive cues amidst illusory cutaneous contact.

## Introduction

We integrate a multimodal array of sensorimotor inputs in the everyday perception of our natural environment. Along with vision and audition, our sense of touch is essential in interactions involving dexterous manipulation, affective connections, and naturalistic exploration (*1*–*4*). For example, we routinely judge the ripeness of fruit at the grocery store, caress the arm of a spouse to offer comfort, and stroke textiles to gauge their roughness and softness (*5, 6*). We seamlessly do so by recruiting sensorimotor inputs, fine-tuning motor control strategies, comparing to prior experiences, and updating internal representations (*7*).

Historically, illusions have revealed inherent interdependencies of our sensorimotor and perceptual systems. Among the many tactile illusions identified (*8, 9*), the “size-weight” illusion is particularly well-known. It involves picking up two objects of identical mass but of varied volume, and indicates that the smaller object is generally perceived as heavier (*10*). The size-weight illusion reveals a separation of our sensorimotor and perceptual systems in estimating an object’s mass. In particular, while our sensorimotor system adapts to the mismatch between the predicted and actual signals to dynamically adjust our exploratory motions, our perceptual system recalibrates the size-weight relationship more gradually on a different time scale (*8, 11*). Another intriguing illusion regards our perception of curvature. With it, a flat physical surface can be perceived by the bare finger as curved, depending on the relative lateral motion between the finger and surface (*12*). The curvature illusion reveals a poor spatial constancy of our somatosensory system, driven by a dissociation between cutaneous and proprioceptive inputs (*3*). This illusion is practically important in rendering haptic interactions in 3D virtual environments (*8*). A further illusion, in analogy with the Aubert-Fleischl phenomenon in vision, and Doppler effect in audition, indicates that the speed of a moving stimulus can be overestimated by touch amidst changes in proprioceptive motion (*13*). These and other perceptual illusions have shed light upon interdependencies of our sensorimotor and perceptual systems, and lend to subsequent engineering applications.

Among the many dimensions of touch, which include surface roughness, stickiness, geometry, and others, our perception of softness is central to everyday life (*2*). Our understanding of tactile compliance, a key dimension of an object’s “softness,” remains incomplete. This percept is informed by some combination of cutaneous inputs from mechanosensitive afferents signaling skin deformation and proprioceptive inputs signaling body movements. Efforts to define the precise cues within skin deformation and body movements have focused on contact area at the finger pad (*14*–*18*), spatiotemporal deformation of the skin’s surface (*19*–*21*), and kinesthetic inputs of displacement, force, and joint angle (*22*–*25*). Such an array of sensory contact inputs, mediated by independent cortical mechanisms, are recruited and integrated in the primary somatosensory cortex, and form the perceptual basis from which compliances are recognized and discriminated (*26*).

Here, we identify a tactile illusion that underlies our perception of softness, or compliance, which is produced by covarying the elasticity and curvature of spherical stimuli. These attributes are routinely encountered, such as in judging the ripeness of fruit. The illusion, however, is only observed in passive touch. The stimuli become readily discriminable when explored volitionally. This illusion therefore naturally decouples cutaneous cues from proprioceptive cues, and sheds light into how we volitionally explore compliant objects.

## Results

We introduce a novel elasticity-curvature illusion where small-compliant and large-stiff spheres are perceived as identical. This is done for single, bare finger touch. Our methodological paradigm is unique in that computational models of the skin’s mechanics define the stimulus attributes prior to evaluation in human-subjects experiments. In particular, finite element models of the distal finger pad are used to develop elasticity-curvature combinations that generate non-differentiable cutaneous contact. Then, validation of the illusion is done empirically in human-subjects via measurements of biomechanical interactions of the finger pad and evaluations of psychophysical responses. The elasticity-curvature illusion suggests that we use a force-controlled movement strategy to optimally evoke cutaneous and proprioceptive cues in discriminating compliance.

First, the skin mechanics of the index finger are modeled with finite elements in simulated interactions with spherical stimuli. The models predict that small-compliant (10 kPa – 4 mm) and large-stiff (90 kPa – 8 mm) spheres will generate nearly identical cutaneous contacts, which may render them indiscriminable in passive touch. In contrast, when the models simulate conditions of active touch, the resultant fingertip displacements with controlled force loads are found to be distinct, which may render them discriminable.

Next, driven by the model predictions, a series of biomechanical and psychophysical evaluations are conducted with human participants. The results reveal that these spheres are indeed indiscriminable when explored in passive touch with only cutaneous cues available. However, this illusion vanishes when cues akin to proprioception are systemically augmented by a participant’s use of a force control movement strategy.

### Experiment 1: Computational modeling of the elasticity-curvature illusion

Finite element analysis was performed to simulate the skin mechanics of the bare finger interacting with compliant stimuli. The material properties of the model were first fitted to known experimental data. Then, numerical simulations were conducted with spherical stimuli of covaried radius (4, 6, and 8 mm) and elasticity (10, 50, and 90 kPa). In two interaction cases, the fingertip was moved and constrained to simulate active and passive touch, respectively. To help quantify the discriminability of the spheres, response variables were derived from the stress distributions at the epidermal-dermal interface, where Merkel cell end-organs of slowly adapting type I afferents and Meissner corpuscles of rapidly adapting afferents reside, as well as the required fingertip displacements to a designated touch force. The former was deemed as the cutaneous cue (*21, 27, 28*). The latter was associated with the proprioceptive cue where displacement approximates the change in muscle length and force tied to muscle tension (*24, 28*–*31*).

In simulation of passive touch where only cutaneous cues are available, the compliant spheres deformed the surface of the skin distinctly for each combination of elasticity and radius (Fig. 1). Spatial distributions of stress for both the finger pad and spheres were simulated to a steady-state load of 2 N. For all the nine spheres simulated, either an increase of the spherical radius or a decrease of the elasticity decreases the concentration of stress quantities at contact locations, with the lowest stress concentration for the10 kPa-8 mm sphere and the highest for the 90 kPa-4 mm sphere (detailed in Fig. S1).

**Fig. 1.**
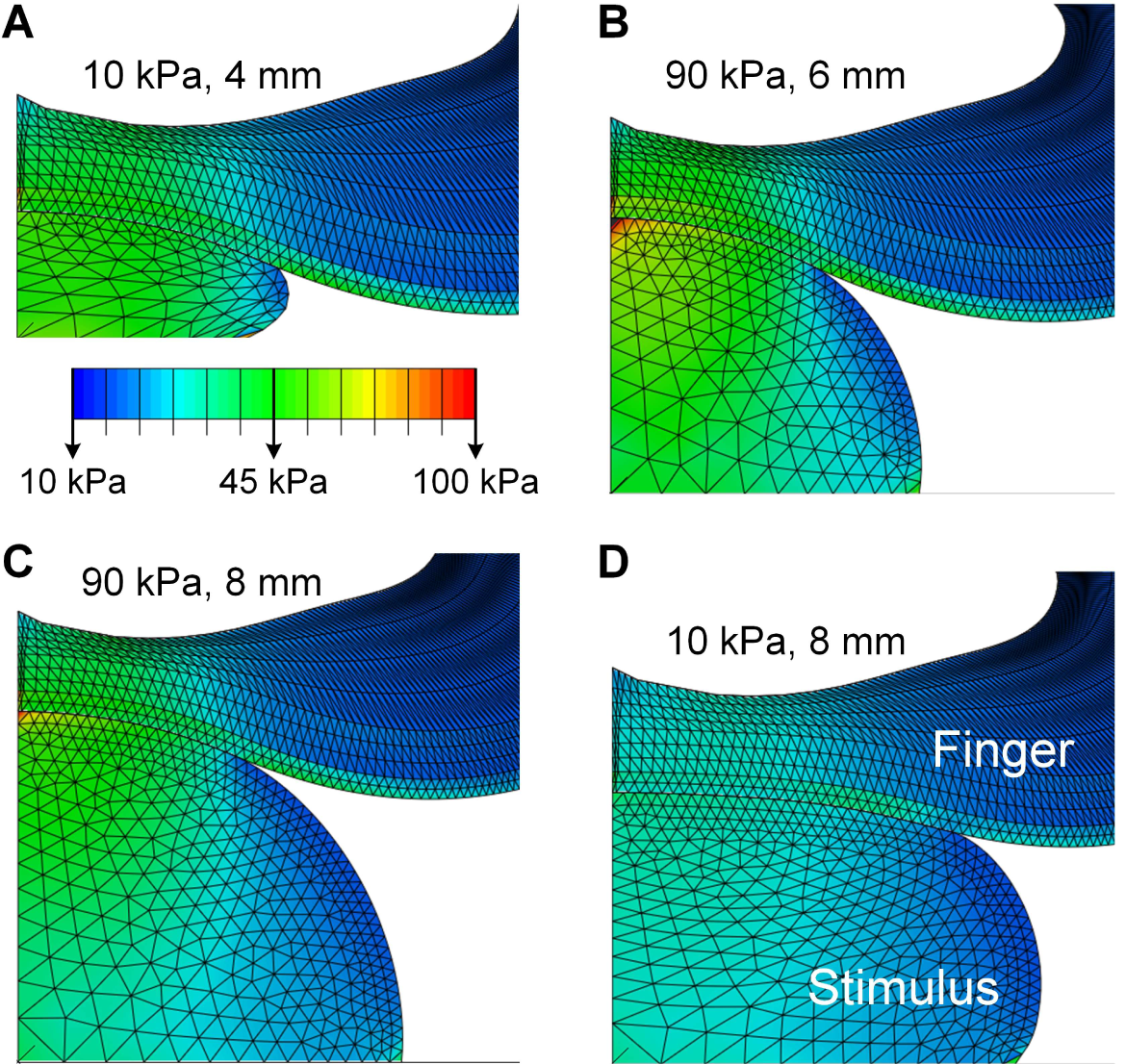
Computational modeling of contact mechanics with compliant spheres. Spatial distributions of stress are simulated at a load of 2 N for contact with spheres of 10 kPa-4 mm (**A**), 90 kPa-6 mm (**B**), 90 kPa-8 mm (**C**), and 10 kPa-8 mm (**D**) respectively. Although the deformation of the spherical stimuli differs greatly from (**A**) to (**C**), the resultant stress distributions and surface deflection at the finger pad are nearly identical.

However, for certain elasticity-radius combinations, changes in the spheres’ radii counteracted the changes in their elasticity, resulting in identical stress distributions for those cutaneous contacts. Although the deformation of the stimuli differed vastly between the 10 kPa-4 mm (Fig. 1A) and 90 kPa-8 mm spheres (Fig. 1C), the surface deformation and stress distributions of the finger pad were quite similar. Specifically, in Fig. 2A, stress distributions at the contact interface were nearly identical between the small-compliant (10 kPa-4 mm) and large-stiff (90 kPa-8 mm) spheres across all levels of load. A similar case is demonstrated for the 10 kPa-4 mm and 90 kPa-6 mm spheres where the stress curves fairly well overlap (Fig. 2B), as compared to a sphere with the same elasticity but distinct radii (Fig. 2C).

**Fig. 2.**
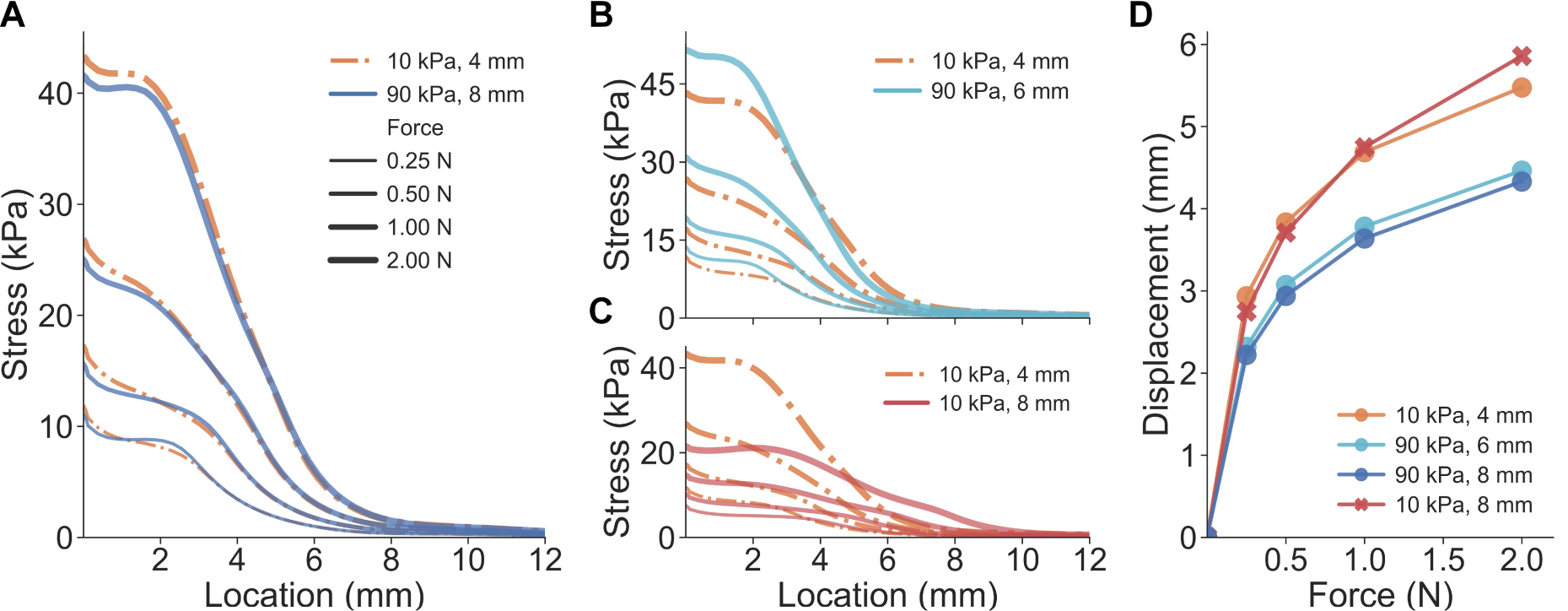
Results of Experiment 1: Cues of cutaneous contact and proprioception. (**A**) For the small-compliant (10 kPa-4 mm) and large-stiff (90 kPa-8 mm) spheres, stress distributions at adjacent locations are nearly identical across all force loads. (**B**) Curves of stress distributions fairly well overlap for the 10 kPa-4 mm and 90 kPa-6 mm spheres. (**C**) Distinct stress distributions were obtained for spheres with the same elasticity but varied radii. (**D**) Proprioceptive cues of finger displacement are simulated for all force loads.

In addition to spatial distributions of stress, other deformation variables were also evaluated. The strain energy density (SED) and the deflection of the skin’s surface were calculated and analyzed (Fig. S1, S2, and S3). Similar to the stress distributions, responses from the three spheres were nearly inseparable. This demonstrates that certain compliant stimuli – e.g., 4 mm spheres of 10 kPa and 8 mm of 90 kPa – generate nearly identical cutaneous cues, and therefore, naturally dissociate from proprioceptive cues. It indicates that in passive touch where only cutaneous cues are available, one might be unable to differentiate the small-compliant and large stiff spheres.

In simulation of active touch, where both cutaneous and proprioceptive cues are available, an increase in either the radius or elasticity decreases the fingertip displacement given the same load (Fig. S3). Specifically, the force-displacement curve of the 10 kPa-4 mm sphere was clearly separable from the 90 kPa-8 mm sphere (Fig. 2D). Additionally, spheres of the same elasticity yielded overlapping force-displacement curves, as opposed to spheres of different elasticity. These results demonstrate that distinct proprioceptive cues tied to fingertip displacement differ given the indentation of the small-compliant compared to the large-stiff spheres. In active touch, where cues tied to fingertip displacement are utilized, one might be able to perceptually discriminate those illusion case spheres amidst non-differentiable cutaneous contact.

### Experiment 2: Biomechanical measurement of cutaneous contact

Derived from the computational analysis in Experiment 1, we hypothesized that non-differentiable cutaneous contact might be observed between the small-compliant and large-stiff spheres. To validate this prediction, we conducted biomechanical measurement experiments with human-subjects.

In particular, through a series of biomechanical measurements, the contact area between the finger pad and stimulus was quantified to determine if the illusion case spheres would generate similar cutaneous contact. Contact area was measured directly, using an ink-based procedure (*20*). Measured contact area is commensurate with the cutaneous cues predicted in the finite element simulation. In the simulation, stress/strain distributions at contact locations and the skin surface deflection are quantified as the cutaneous cues. In the experiments, contact area is derived from a contiguous area on the skin surface with a super-threshold contact pressure (*16, 17*). Furthermore, the deflection of the skin surface is the contour of a deflection profile in the contact plane (*21, 28*).

In passive touch, where compliant stimuli are indented into a fixed fingertip, a customized indenter was utilized (Fig. 3A). Participants (*n* = 10) were instructed to rest their forearm and wrist on a stationary armrest and the index finger was constrained. Each sphere was indented into the finger pad with a triangle-wave force profile peaking at the desired level (1, 2, and 3 N). To quantify the contact area at the peak magnitude of indentation, an ink-based procedure was employed. The stamped finger pad was digitized (Fig. 3C) and the contact region was color-enhanced (Fig. 3D). The contact areas were then calculated based on the exterior outlines with scaled pixels (Fig. 3E).

**Fig. 3.**
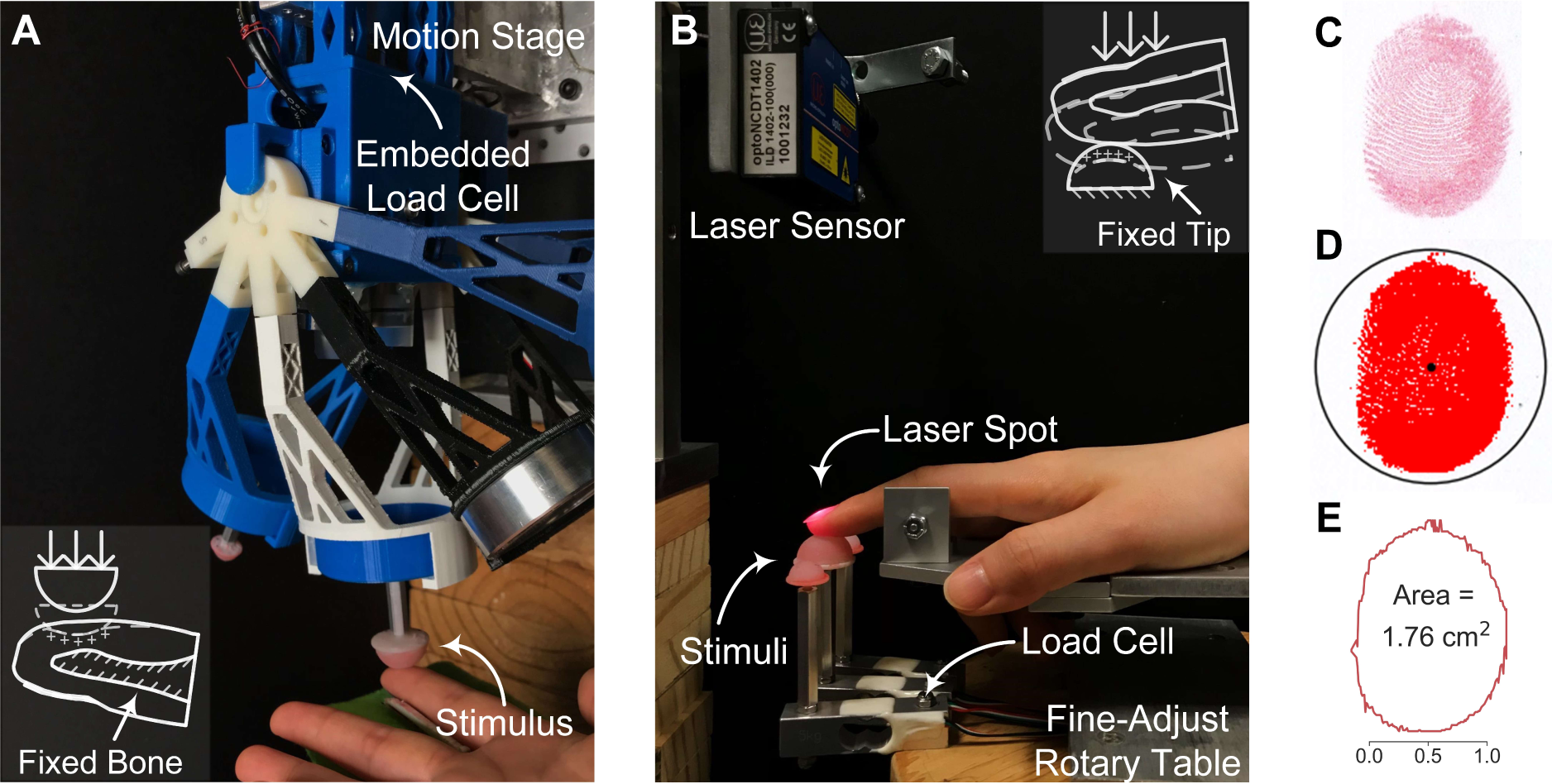
Experimental setup and ink-based contact area analysis. (**A**) For passive touch, the compliant stimulus is indented into the fixed finger pad by the motion stage. Contact force is measured by the embedded load cell. (**B**) For active touch, the designated stimulus is fixed and volitionally contacted by the index finger. Touch force is measured by the load cell underneath and fingertip displacement is captured by the laser sensor. (**C**) Contacted fingerprints are stamped and digitized for analysis. (**D**) The contact region is identified and color-thresholded. (**E**) Contact area is calculated based on the exterior outline and scaled pixels.

Non-distinct relationships of touch force and contact area are indeed observed in passive touch between the illusion case spheres across loading levels (Fig. 4). By inspecting individual trials (Fig. 4A), illusion case spheres (small-compliant and large-stiff) generated similar contact areas while the distinct sphere (10 kPa-8 mm) afforded significantly higher contact areas (*U* = 0.0, *p* < 0.0001, *d* = 4.81). In particular, data points for the three illusion cases were well clustered together across all force levels (0.90 ± 0.12 cm^2^, mean ± SD), while the others were well separated (1.68 ± 0.18 cm^2^). For all participants aggregated (Fig. 4C), the force-contact area relation appeared to be consistent within an individual. Traces for the three illusion cases well overlapped (no significant difference detected) across all force levels, while the trace for the 10 kPa-8 mm sphere was significantly distinct (*U* = 87.0, *p* < 0.0001, *d* = 4.49).

**Fig. 4.**
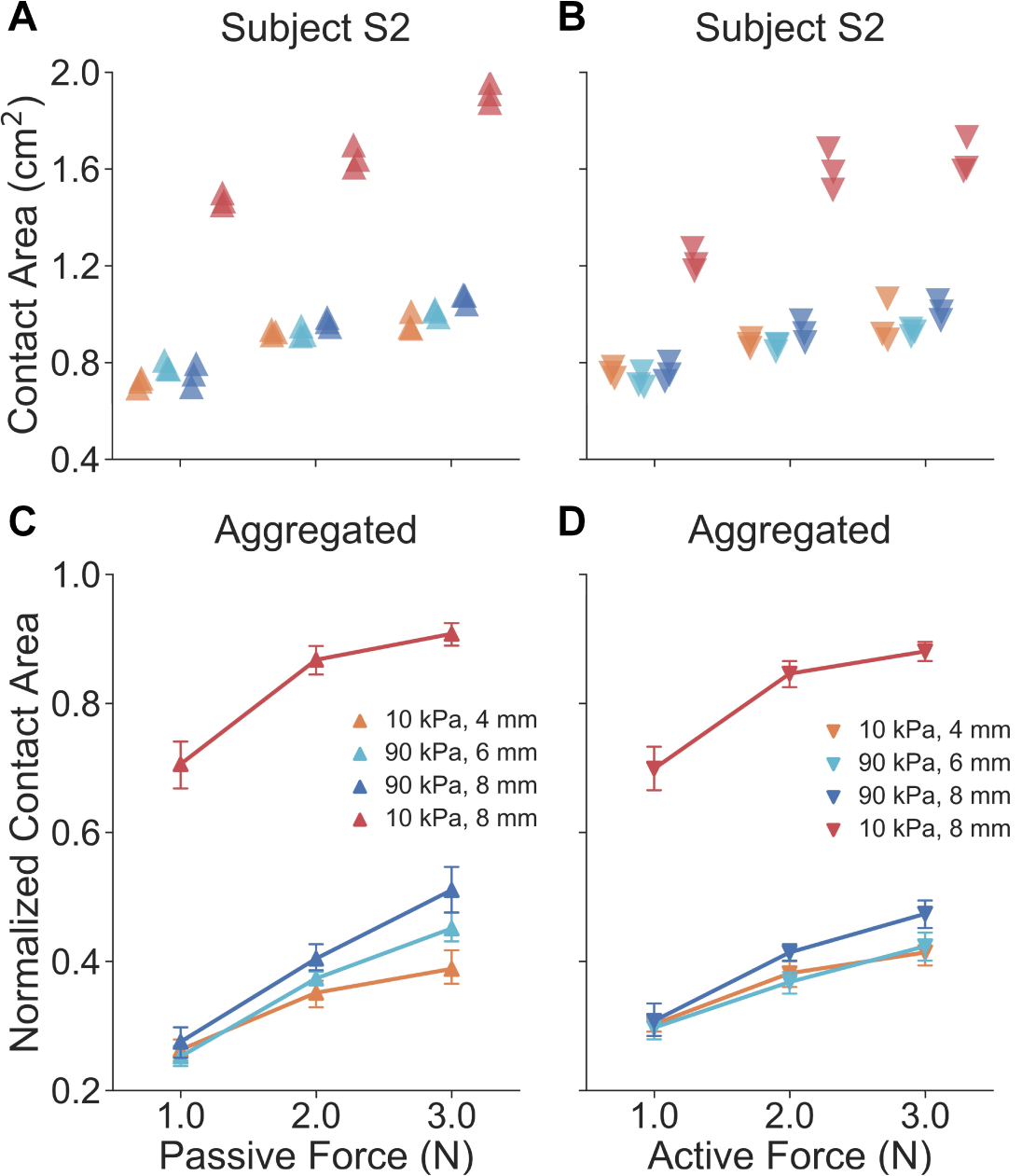
Results of Experiment 2: Biomechanical measurements of contact area. For a representative participant, in both passive (**A**) and active (**B**) touch, gross contact areas for illusion case spheres across all force levels are nearly identical, as opposed to the 10 kPa-8 mm sphere. For all participants aggregated, both in passive (**C**) and active (**D**) touch, curves of the illusion cases well overlap across all force levels, as opposed to the 10 kPa-8 mm sphere. Error bars denote 95% confidence intervals.

In active touch, where the finger volitionally touches the fixed compliant stimulus, an experimental setup was built as illustrated in Fig. 3B. Participants (*n* = 10) were instructed to press their index finger down into a spherical stimulus without external constraint. A sound alarm was triggered to end each trial when the touch force reached the desired level. After each trial, the ink-based procedure was conducted to measure the contact area between the finger pad and stimulus.

Similar force-contact area relations were found in active touch as likewise found in passive touch. Within a participant (Fig. 4B), and similar to the passive touch experiments, the illusion case spheres indeed generated identical gross contact areas while the 10 kPa-8 mm sphere exhibited significantly distinct results (*U* = 0.0, *p* < 0.0001, *d* = 3.73). Specifically, data points for the illusion cases were clustered around 0.87 ± 0.10 cm^2^ while the others were quite distinct from 1.48 ± 0.20 cm^2^ across all force levels. For all participants aggregated (Fig. 4D), traces for the illusion cases well overlapped (no significant difference detected), and the 10 kPa-8 mm sphere yields a much more distinct relationship (*U* = 0.0, *p* < 0.0001, *d* = 4.49). Because cues tied to contact area are identical, proprioceptive inputs evoked in active touch may be vital to discriminating the illusion case spheres.

### Experiment 3: Psychophysical evaluation of the elasticity-curvature illusion

The results of Experiment 2 support the hypothesis that cutaneous contact is indiscriminable between small-compliant and large-stiff spheres, for both passive and active touch. To validate whether these compliant spheres might indeed render a perceptual illusion, we conducted psychophysical experiments with human-subjects.

To test this hypothesis, participants (*n* = 10) were first instructed to discriminate the illusion case spheres in passive touch. In the “passive same force-rate” task, where indentation rate was controlled (Fig. 5A), participants were not able to discriminate the stimuli (detection rate of 46.1% ± 5.7). This illustrates that, when only cutaneous cues are available for discrimination but their contact areas found non-differentiable, these spheres indeed generate a perceptual illusion.

**Fig. 5.**
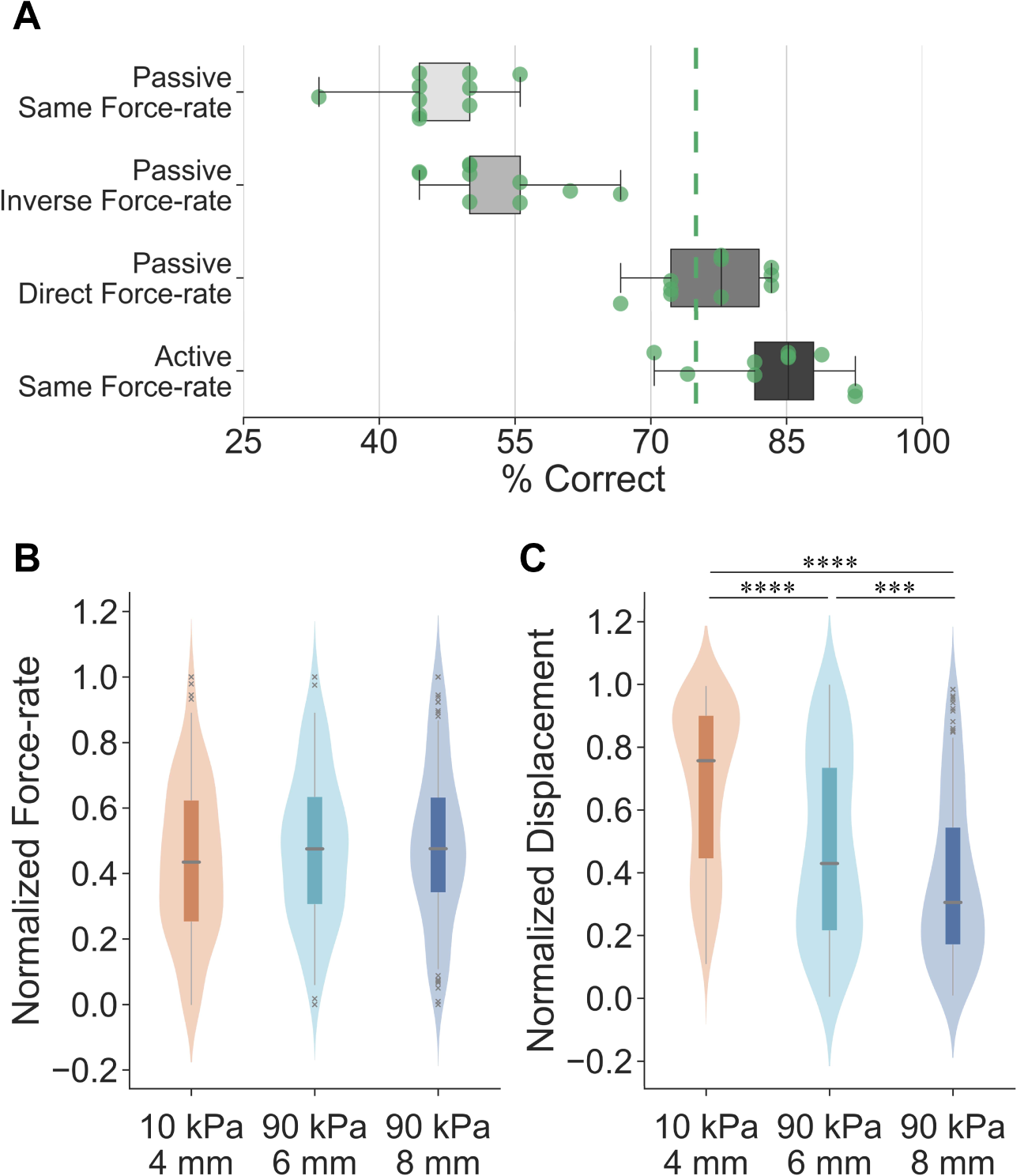
Results of Experiment 3: Psychophysical evaluations and exploratory strategies. **(A)** Psychophysical evaluations of illusion case spheres under different experimental conditions with all participants aggregated. The detection threshold is set as 75% for the same-different procedure. Points denote individual results. (**B**) Non-distinct force-rate cues are volitionally applied for each illusion case sphere in active exploration of compliances. A miniature boxplot is set in the interior of the kernel density estimation of the underlying distribution. (**C**) Significantly higher fingertip displacement is applied for the small-compliant sphere, as opposed to the harder spheres. \*\*\**p* < 0.001, \*\*\*\**p* < 0.0001.

Then, to evaluate if the elasticity-curvature illusion becomes discriminable when adding proprioception to cutaneous contact, controlled force inputs were induced in passive touch in two separate cases. In the “passive inverse force-rate” task (Fig. 5A), where the softer stimulus was indented “inversely” at a higher force-rate than the harder stimulus, participants were still unable to discriminate the compliances with a detection rate of 52.8% ± 6.7. However, this performance improved significantly as compared with “passive same force-rate” (*U* = 24.0, *p* < 0.05, *d* = 1.03). In the “passive direct force-rate” task, where the softer stimulus was indented “directly” at a lower force-rate than the harder stimulus, participants could differentiate the illusion near a 75% threshold (76.7% ± 5.4). This performance is significantly improved compared to the “passive inverse force-rate” task (*U* = 0.5, *p* < 0.0001, *d* = 3.72). These results empirically validate that when cues additional to those cutaneous are available, the illusion case spheres become discriminable. It further indicates that the controlled force-rate cues are likely perceived akin to proprioception and vital for discrimination, a point which will be detailed in the Discussion.

Fourth, to validate the hypothesis that the proprioceptive cue of active finger displacement may help to discriminate the illusion, psychophysical evaluations were conducted in active touch. Participants (*n* = 10) were instructed to discriminate the illusion case spheres under fully active, volitional sensorimotor control. As illustrated in Fig. 5A, the illusion was readily discriminable with a detection rate of 83.7% ± 6.9 (see Table. S2 for detailed results). This presents significantly improved performance compared to the “passive direct force-rate” task (*U* = 21.0, *p* < 0.05, *d* = 1.08). Altogether, the proprioceptive cues elicited by active, volitional control of finger movements, help in discriminating the stimuli amidst illusory cutaneous contact.

Furthermore, in active touch, participants volitionally move their fingers to generate consistent force trajectories between stimuli (Fig. 5B) and thereby utilize the resultant differences in the fingertip displacements between the illusion stimuli to discriminate them (Fig. 5C). Specifically, given the same force-related cues among illusion cases, significantly higher displacement was applied for the softer spheres (*U* = 8786.0, *p* < 0.0001, *d* = 0.74; *U* = 5737.5, *p* < 0.0001, *d* = 1.18). This finding aligns with the finite element simulation where the 10 kPa-4 mm sphere exhibited higher fingertip displacement under the same force load (Fig. 2D). In summary, when cutaneous cues as well as force-related movement cues are controlled, elicited differences in fingertip displacements help discriminate the illusion case spheres.

## Discussion

This study identifies an elasticity-curvature illusion underlying our perception of compliance. The two attributes of this perceptual illusion are common to everyday naturalistic tasks, for example, in judging the ripeness of fruit. Through a combination of solid mechanics modeling, biomechanical contact measurement, and psychophysical evaluation, we show that small-compliant and large-stiff spheres generate similar cutaneous contact, which give the illusion that two very different objects are the same. However, this illusion vanishes in active touch, when cues akin to proprioception augment indiscriminable cutaneous contact. Furthermore, the results indicate that in the exploration of compliant objects, force-controlled movements are more efficient and optimal for eliciting the cutaneous and proprioceptive cues that underlie our judgments of compliance.

### A force-control movement strategy is optimal, efficient, and underlies softness perception

Amidst illusory cutaneous contact, participants’ volitionally control the contact forces they apply to soft objects. Such force-modulated movement is efficient and optimal at evoking differences in fingertip displacement. These fingertip displacements are proprioceptive by nature and critical to the discrimination of the illusory stimuli (Fig. 5B and C). Indeed, this exploratory strategy is important from a number of other perspectives. First, a force-modulation strategy is essential to compensate for the natural remodeling of the skin over time, which leads to changes in its thickness and elasticity (*21*). Such changes in the skin’s mechanics could generate large variance in neural firing patterns, and thereby perception. However, the skin can reliably convey information about indentation magnitude, rate, and spatial geometry when touch interactions are controlled by surface pressure. Since force directly converts to pressure on the skin upon contact, a force-modulation strategy echoes theories of active, volitional control when exploring soft objects in daily tasks (*21, 23*). Second, at the behavioral level, we prioritize exploratory force to optimize our perception of object compliances in relevant contexts (*23, 29, 32, 33*). Indeed, the availability of force-related cues improve discriminability by reducing the necessary deformation of the skin (*20*). Similarly, for the exploratory procedure of pinch grasp, we control the grip force within a safety margin, informed by skin mechanoreceptors, to prevent slipping or overly grasped (*34*).

### Change of cutaneous contact as a cue to proprioception

As just discussed, the force-control movement strategy is efficient and optimal in evoking differentiable cues, in active touch. In passive touch, we observe that participants can discriminate the illusory stimuli, in the “passive direct force-rate” case, at a rate of about 77% correct performance (Fig. 5A). While lower than the rate for active touch, this represents a significant improvement over the “passive same force-rate” case, which yields chance performance.

We hypothesize that the modulation of force under “passive direct force-rate” – where the softer stimulus was indented “directly” at a lower force-rate than the harder stimulus – provides an alternate perceptible input, also tied to finger proprioception. In particular, in alignment with prior findings (*5, 20*), we show that the rate of change of force is linearly correlated with the rate of change of gross contact area (Fig. S6, see Supplementary Materials). While we cannot directly measure the rate of change of contact area, due to the limitations of the ink-based method only being able to measure terminal contact area, such cues might induce the illusion of fingertip displacement (*30, 35*). In particular, Moscatelli, *et al*. demonstrated that skin deformation of this kind naturally induces a sensation of relative finger displacement to the stationary hand (*14, 36*). Similarly, stretching the skin at the proximal interphalangeal joint can induce illusions of self-motion in anesthetized fingers (*35*). Moreover, microscopic oscillatory stimulation at the skin surface also can elicit illusory finger displacements when pressing on a stiff surface (*37*). Therefore, when passively exploring the illusion case spheres, the improved discriminability is likely derived from proprioceptive sensation of this kind.

Indeed, across a range of touch interactions broader than just softness, we find that cutaneous and proprioceptive cues are integrated to achieve high levels of performance (*3, 26*). In tasks involving reaching movements, cutaneous cues could systematically bias motion estimates, indicating that multisensory cues are optimally integrated for our motor control (*3*). In general, multimodal interactions between these two signals are found to be mediated by distinct neural mechanism in primary somatosensory cortex (*26*). This comes in general agreement with prior studies reporting that both cutaneous and proprioceptive cues are needed in discriminating compliance. In particular, when finger movements are eliminated, our ability to discriminate pairs of spring cells decrease (*22*). Likewise, when pinching an elastic substrate in-between two rigid plates, relatively lower discriminability of compliance is obtained when relying upon proprioception alone as compared to cutaneous cues alone (*19*).

### A naturalistic illusion that underlies everyday tasks

The attributes of elasticity and curvature are found in everyday, ecologically relevant tasks. In some prior studies, however, stimuli have been highly engineered and delivered by sophisticated devices (*9*). Such stimuli may not afford the same perceptual acuity as ecologically accurate soft objects (*6*). Moreover, stimulus compliance at times has been parameterized by its stiffness rather than its modulus (*19, 22, 28*), which can be confounding for naturalistic objects of identical stiffness but differing in geometry (*28*). Furthermore, stimuli with flat surfaces do not fully mimic the contact profile of the skin surface’s contacting an elastic object (*8, 9, 16*). Herein, we address these issues by building spherical stimuli with combinations of radii and elasticity which can recapitulate properties of ecologically compliant materials, such as fruit. This enables us to identify a perceptual illusion that is naturalistic and commonplace as our daily tasks. As it is difficult to measure the material properties of fruit, which can breakdown rapidly between sessions, our group has begun to consider the perceptual commonality between silicone-elastomer materials as reasonable stand-ins for ecological fruits (*5*). Similar to the work with engineered substrates herein, we have found that the exploratory strategy of volitionally controlling force aligns with how we judge the ripeness of fruit. In particular, we volitionally pinch soft fruit, by controlling grip force, to help differentiate their ripeness (*5*).

### Computational modeling formulates psychophysical studies

Instead of evaluating empirically with human-subjects a large number of stimulus combinations of elasticity and curvature, we computationally identified combinations with indistinct cutaneous contact. Indeed, a “computation first” effort as such demonstrates an alternative paradigm to bridge theoretical and empirical studies, make specific predictions and test particular hypotheses. Specifically, to better understand the encoding mechanism underlying the identified tactile illusion, cutaneous and proprioceptive cues need to be dissociated. As this is empirically demanding, we employed two interaction modes (passive and active touch, Fig. 3) in the computational simulation. The potential cues and interaction modes that modulate the illusion are then validated in psychophysical experiments with human-subjects.

Finally and relatedly, far fewer illusions have been discussed in the tactile modality than for vision and audition (*8, 9*). This partially reflects the fact that tactile illusions are not as easily accessible (*8*). Indeed, sophisticated efforts are usually required to create appropriate conditions to conceive the illusion, which is a significant electromechanical challenge to achieve empirically (*9*). The “computation first” approach demonstrated herein may help in identifying potential illusions in a more efficient manner.

## Materials and Methods

### Geometry of the fingertip model

Two simplified 2D finite element models were derived from the geometry of a 3D model of the human distal phalange (*28*). The plane strain model of a cross-sectional slice from proximal first digit to distal tip was built for contact across the finger width (Fig. S4). Meanwhile, the axisymmetric model revolving around the centerline of the finger pad was built for contact normal to the surface (Fig. S4). Details of the model’s structure and mesh are further explained in Supplementary Materials.

### Material properties of the fingertip model

Hyperelastic material properties were used of the Neo-Hookean form of the strain energy function. The strain energy Ψ was derived as:

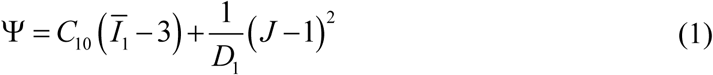

where *C*_10_, *D*_1_ were material constants (*28*), 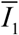 was the modified first strain invariant, and *J* was the volume ratio known as the Jacobian matrix. The initial shear modulus *G* was predefined and the initial bulk modulus was as *K* = *G* /10^5^. The relationship between modulus and material constants were defined as *G* = 2*C*_10_ and *K* = 2 *D*_1_ accordingly.

The material elasticity was defined by its initial shear modulus *G* which fully justified the material. Note that the material is in fact non-linearly hyperelastic. The Neo-Hookean model was applied to simplify the fitting procedure and derive a more robust calibration only based on the modulus *G*. Furthermore, instead of a linear Young’s modulus, the hyperelastic form was considered for the soft objects which deform in a finite-strain region.

Finally, material calibration was conducted in two steps. First, the ratios of material elasticity between each layer were fitted to match the observed surface deflection to different displacements (Fig. S5A). Second, the fitted ratios were scaled to fit the observed force-displacement relationships (Fig. S5B). The detailed fitting procedures and final results are explained in Supplementary Materials.

### Stimulus tip model

Three values of radii (4, 6, and 8 mm) and elasticity (10, 50, and 90 kPa) were selected and the stimulus tips were modeled as hemispherical with the surface of central section attached to a rigid plate. The Poisson’s ratio to the plate was set to 0.475 to mimic the nearly incompressible behavior of rubber. For the purpose of suppressing stress concentrations near nodes, triangular elements with 0.25 mm edge length were used in the region contacting the finger surface. Larger elements of up to 1.0 mm were used in non-contact region to lower the computational cost.

### Numerical simulations

Nine stimulus tips (3 radii by 3 elasticity) were built based on the 2D axisymmetric model and contact mechanics were simulated in attempt to approximate passive and active touch interactions. In passive touch (Fig. 3A), compliant stimuli were indented into the fixed fingertip at loads of 0.25, 0.5, 1, and 2 N. The response variables were derived as cutaneous cues only, quantified by stress distributions at the epidermal-dermal interface (470 μm beneath the skin surface), calculated by averaging neighboring elements at each interface node. Note that proprioceptive cues were decoupled since the reaction force was provided by the fixture instead of muscle activity. In active touch (Fig. 3B), the fingertip was ramped into the fixed stimuli to the aforementioned loads. The response variables were derived from both the cutaneous and proprioceptive cues. Specifically, the proprioceptive cue was approximated by the force-displacement relation of the fingertip in the normal direction. This measure is tied to the change of muscle length as detected by muscle spindles, while force indicates the change of the muscle tension of Golgi tendons (*3, 14, 30*).

### Stimuli and experimental apparatus

The compliant stimuli were constructed from a room temperature curing silicone elastomer (BJB Enterprises, Tustin, CA; TC-5005 A/B/C). To achieve the desired modulus, based on prior calibrations (*16*), corresponding ratios of cross-linker were added and mixed. These formulations were then cast into 3-D printed molds of three radii (4, 6, and 8 mm) and cured to become stimulus tips.

As illustrated in Fig. 3A, a customized motion stage (ILS-100 MVTP, Newport, Irvine, CA) was built to indent the stimulus into the stationary finger pad (*20*). Normal contact force was recorded with a load cell (22.2 N, 300 Hz, LCFD-5, Omega, Sunbury, OH) mounted onto the cantilever. The 3D printed housing fixture was equipped with a servo motor (Parallax standard servo, Rocklin, CA) and actuator arms, enabling a quick switch between different stimuli. Customized circuitry and software were developed to command the indentations. The participant’s forearm was fixed on a stationary armrest with their index finger fixed at approximately 30 degrees relative to the stimulus surface.

The experimental setup for active touch is shown in Fig. 3B. Instrumented load cells (5 kg, 80Hz, TAL220B, HTC Sensor, China) were installed on a fine-adjust rotary table which can be rapidly rotated to present the designated stimulus. To measure the fingertip displacement, a laser triangulation displacement sensor (10 µm, 1.5kHz, optoNCDT ILD 1402-100, Micro-Epsilon, Raleigh, NC) was mounted and the laser beam was calibrated to aim at the center of the stimulus surface. The forearm, wrist, and palm base rested on a parallel beam with no external constrains.

### Measurement of contact area

The gross contact area between the stimulus surface and finger pad was measured by the ink-based method (*16, 20*). An overview of this method is shown in Fig. 3 and summarized as follows. At the beginning of each measurement, washable ink (Craft Smart, Michaels Stores, Inc., Irving, TX) was fully applied onto the stimulus surface. After each contact, the participant was instructed to gently indent the finger pad onto a blank section of a sheet of white paper, to fully transfer the stamped ink. The remaining ink on the finger pad was then completely removed. This procedure was repeated until all trials were completed for the participant. The sheet of paper was then marked with a 5.0 cm reference bar and digitized for analysis. A center-radius pair was selected by the analyst to identify a region enclosing the fingerprint. The desired color rendering was adjusted to outline the edges from the background. Next, a serial search was conducted to find these bounding edges and the reference bar was also identified to scale the pixels. The final area was calculated using Gauss’s formula in squared centimeters.

### Measurement of force and displacement

Readings from the force and laser sensor were smoothed to remove electrical artifacts by a moving filter with a window of 100 neighboring readings. The ramp segments of the force curves were then extracted based on first-order derivatives (*5*). A linear regression was applied to the segments and the derived slope was noted as the force-rate. On the other hand, the fingertip displacement was calculated as the absolute difference between the initiation and conclusion of each movement.

### Participants

The human-subjects experiments were approved by the Institutional Review Board at the University of Virginia. Ten naïve participants were recruited (5 females and 5 males, 27.5 ± 2.6 years of age) and provided informed consent. No history of upper extremity pathology that might impact sensorimotor function was reported. All participants were right-handed and were assigned to complete both the biomechanical and psychophysical experiments. All experimental tasks were completed and no data were discarded.

### Experiment procedure

In Experiment 2, the biomechanical measurement experiments were conducted in both passive and active touch. For passive touch, the four stimuli were each indented into the finger pad at three force levels (1, 2, and 3 N) during three sessions respectively. Each stimulus was ramped into the finger pad for one second and retracted away for one second. The ink-based procedure was applied for each indentation. For each participant, there were three trials for each stimulus for each indentation level. All trials were separated by a 20-second break. For active touch, the four stimuli were palpated by the index finger at the same three force levels which were behaviorally controlled and presented during three sessions respectively. In particular, participants were instructed to actively press into the designated stimulus and a sound alarm was triggered to end the current trial when their force reached the desired level. The ink-based procedure was used for each trial. There were three trials for each stimulus for each force level for each participant. All trials were separated by a 20-second break.

In Experiment 3, psychophysical discrimination experiments were conducted for both passive and active touch. Following the rule of ordered sampling with replacement, nine stimulus pairs were drawn from the three illusion cases and were prepared for psychophysical evaluation. Participants were blindfolded to eliminate any visual information about the stimulus compliance or the movements of the indenter and the finger pad. No feedback on their performance was provided during the experiment. Using the same-different procedure, after exploring each pair (one touch per stimulus), participants were instructed to report whether the compliances of the two were the same or different. Note that the same-different procedure was applied herein because the observer can use whatever cues are available and does not have to articulate the ways in which the compliances actually differ (*38, 39*). This fits well with the experimental scope where the roles of perceptual cues are under investigation.

For passive touch, within each trial for a pair of stimuli, spheres from the same pair were ramped into the fixed finger pad successively (Fig. 3A). The indentation interval was controlled as 2-seconds to obtain consistent temporal effects on perception (*40*). Each of the nine stimulus pairs was presented twice per each participant. The test order of all trials was randomized to balance the effects of fatigue or inattention. All discrimination trials were separated by a 15-second break. The terminal force level was set to 2 N as this aligned with Experiments 1 and 2. As illustrated in Fig. 5, three experimental tasks were performed in passive touch. In the “passive same force-rate” task, all stimuli were indented at 1 N/s to 2 N. In the “passive inverse force-rate” task, higher force-rate was applied for the soft stimulus while the lower force-rate was applied for the hard stimulus. The 10 kPa-4 mm sphere was indented at 2 N/s to 2 N. The 90 kPa-6 mm and 90 kPa-8 mm sphere were indented at 0.5 N/s to 2 N. In the “passive direct force-rate” task, force-rate was applied in a direct positive relation with the stimulus modulus. The 10 kPa-4 mm sphere was indented at 0.5 N/s to 2 N and the two 90 kPa spheres were indented at 2 N/s to 2 N.

For active touch, the experiments were conducted under participants’ fully active, behavioral control (Fig. 3B). In particular, a participant was instructed to explore compliance by palpating each of two spheres successively. When their touch force reached 2 N, a sound alarm was triggered to end that exploration. The interval between two trials was set to 2-seconds as previously noted. Force and fingertip displacement were recorded simultaneously. Each stimulus pair was presented twice in a randomized order and there was a 15-seconds break between every two trials.

### Data analysis

As illustrated in Figs. 4 and 5, the experimental results for all participants were aggregated for analysis. A normalization procedure was required for data aggregation since participants exhibited distinct sensorimotor capabilities, range of finger movements, and dimensions of the finger pad (*5, 20*). In particular, for each experimental task per each participant, all recordings of each tactile cue were normalized to the range of (0, 1) by sigmoidal membership function (*5, 24*). The center of the transition area was set as the average value of the data normalized, and the logistic growth rate of the curve was set to 1. After this transition, data from all participants were aggregated together for statistical analysis. The Mann–Whitney U test (*α* = 0.05, two-sided test) was applied to compare the sample means and the Cohen’s *d* (the absolute value) was calculated for statistically significant results to evaluate the effect size. The confidence interval was derived by bootstrapping the estimated data with 1000 iterations.

## Supplementary Materials

Perceptual cues predicted in the computational modeling

Geometry of the fingertip model

Fitting hyperelastic material properties

Perceptual cues measured from one representative participant

Fig. S1. Simulated spatial distributions of cutaneous cues.

Fig. S2. Comparison of cutaneous cues between illusion and distinct spheres.

Fig. S3. Cues of the surface deflection and finger displacement.

Fig. S4. Geometry of the finger and stimulus tip model.

Fig. S5. Results of the material properties fitting.

Fig. S6. Perceptual cues measured in human-subjects experiments.

Table S1. Material properties derived from the fitting.

Table S2. Results of psychophysical evaluations for all stimulus pairs.

## General

We thank all the participants of the human-subjects experiments.

## Funding

This work is supported in part by grants from the National Science Foundation (IIS-1908115) and National Institutes of Health (NINDS R01NS105241).

## Author contributions

C.X., Y.W., and G.J.G. conceptualized and designed the study. C.X., Y.W., and G.J.G. developed the computational model. C.X., Y.W., and G.J.G. implemented the apparatus and performed the experiments. All authors analyzed and interpreted experimental results. All authors edited and approved the manuscript.

## Competing interests

The authors declare no conflict interests.

## Data and materials availability

Datasets for the current study are available from the corresponding author upon reasonable request. All data required to evaluate the conclusion are included in the paper and/or the Supplementary Materials.

## Supplementary Materials

### Perceptual cues predicted in the computational modeling

For the cutaneous-only cues, besides spatial distributions of stress, other response variables were also derived from the finger-stimulus contact mechanics. Strain energy density (SED) at the epidermal-dermal interface where Merkel cell end-organs of slowly adapting type I (SAI) afferents reside were estimated by averaging neighboring elements at each interface node. As illustrated in Fig. S1, similar distributions were obtained for the cue of stress/strain across all spheres. Either an increase of the elasticity or a decrease of the sphere radius will increase the concentration of stress/strain at contact locations, with the lowest for the 10 kPa-8 mm sphere and the highest for the 90 kPa-4 mm sphere.

Furthermore, the variations in radius counteracted the changes in elasticity, resulting in nearly identical stress/strain distributions at the same contact locations. As illustrated in Fig. S2A-B, compared with the distinct combination of 10 kPa-4/8 mm, overlapping curves were obtained for small-compliant (10 kPa-4 mm) and large-stiff spheres (90 kPa-8 mm). In addition, curves for the 10 kPa-4 mm and 90 kPa-6 mm spheres were fairly similar. These results were consistent with spatial distributions of stress shown in Fig. 2.

**Fig. S1.**
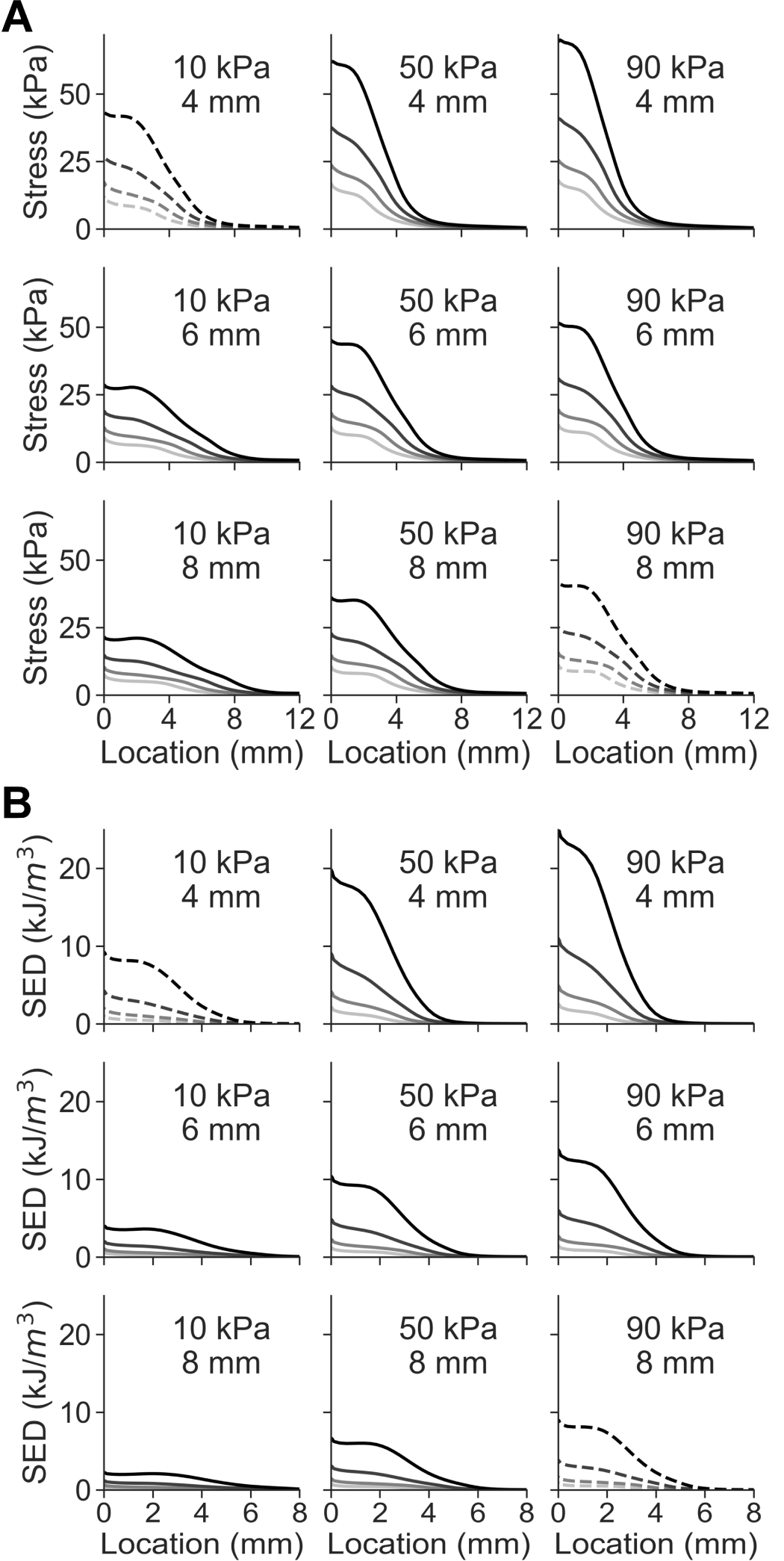
Simulated spatial distributions of cutaneous cues. (**A**) Spatial distributions of stress at contact locations for all nine spherical stimuli. (**B**) Spatial distributions of SED at the same contact locations for all spheres varying in radii and elasticity.

Besides the stress/strain cue at the locations of mechanoreceptive end organs, deflection of the skin surface is often considered as an cutaneous cue informing the change of contact area (*14, 16*). Specifically, deflection of the skin’s surface is the contour of a deflection profile in the contact plane, readily obtainable by visual observation through sophisticated cameras (*17, 41*).

**Fig. S2.**
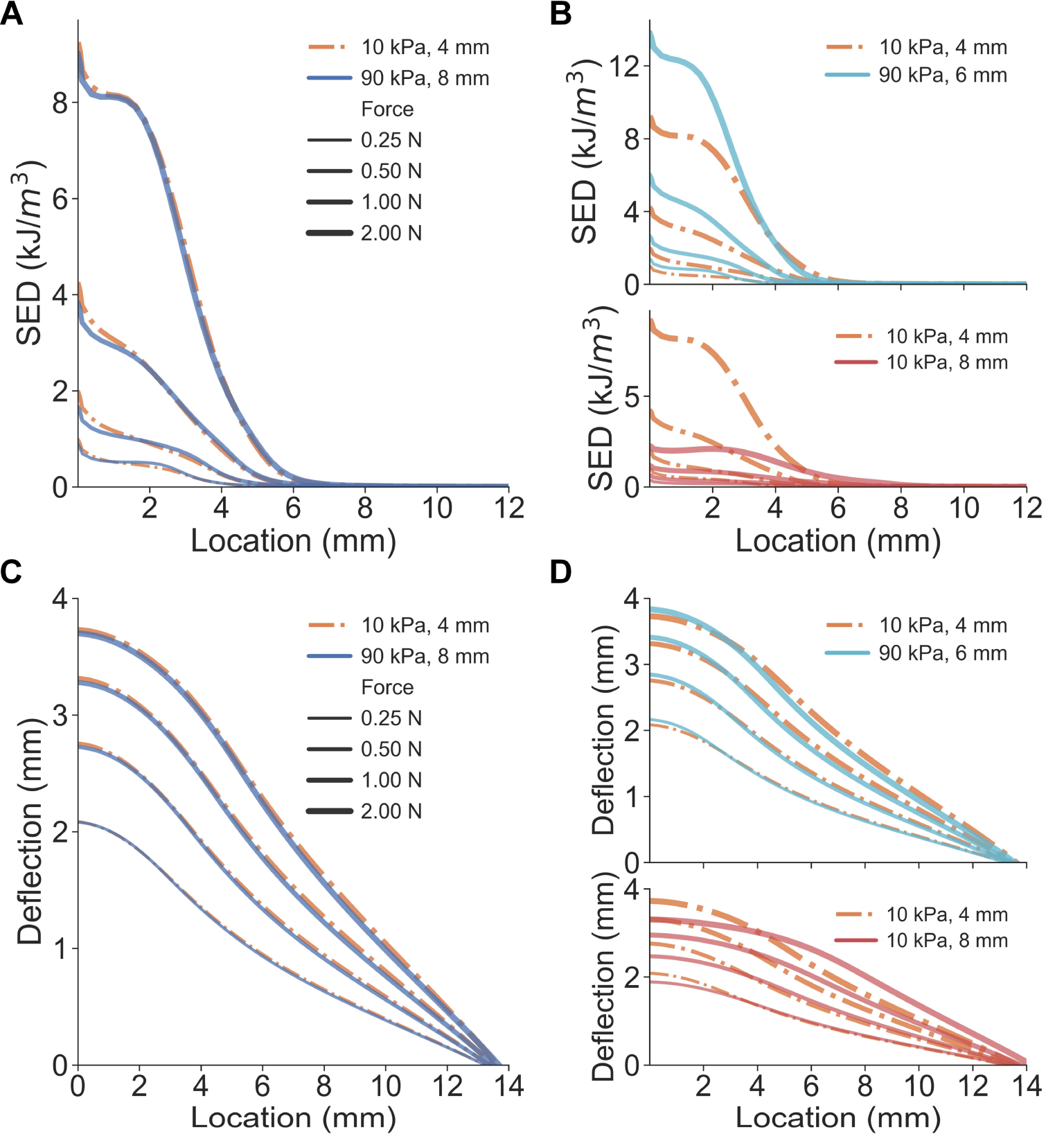
Comparison of cutaneous cues between illusion and distinct spheres. (**A**) Spatial distributions of SED are nearly identical for the illusion case spheres. (**B**) As opposed to the 10 kPa-8 mm sphere, SED distributions fairly well overlap between the 10 kPa-4 mm and 90 kPa-6 mm spheres. (**C**) Non-distinct surface deflection cues are obtained from the illusion case spheres. (**D**) Consistent with SED distributions, surface deflections overlap for the 10 kPa-4 mm and 90 kPa-6 mm spheres.

Therefore, displacements at the node of the epidermis surface were calculated as the skin deflection cue. Similar to results of stress/strain distributions, overlapping curves were obtained from the same stimuli pair (i.e., 10 kPa-4 mm and 90 kPa-8 mm, Fig. S2C). Additionally, as in Fig. S2D, deflection curves for the 10 kPa-4 mm and 90 kPa-6 mm spheres were inseparable, which were predicted to generate nearly identical contact area cues. Overall, as illustrated in Fig. S3A, an increase in the elasticity or a decrease in the spherical radius will increase the magnitude of the surface deflection. Consistent trends were obtained for stress and SED distributions (Fig. S1).

**Fig. S3.**
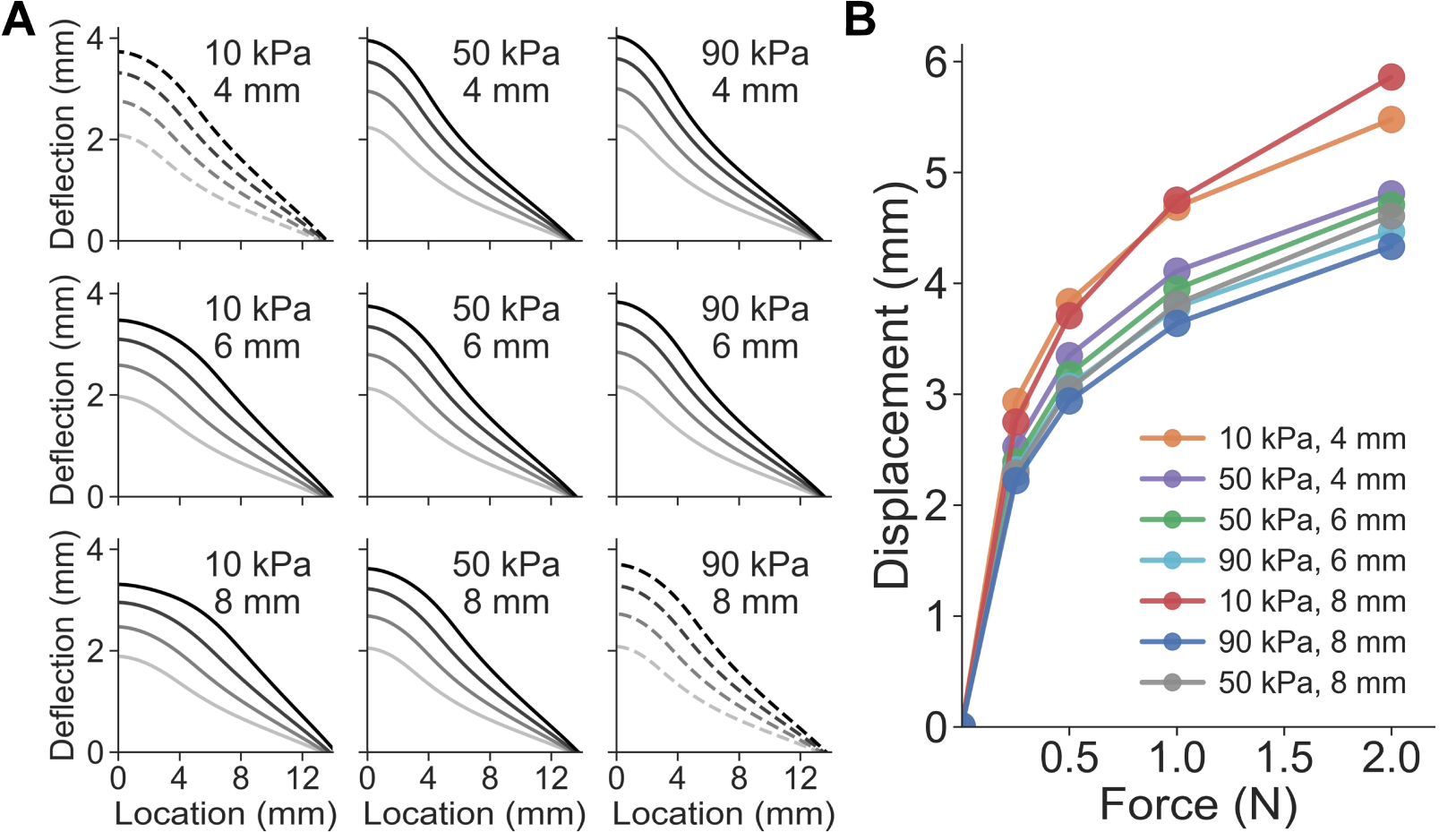
Cues of the surface deflection and finger displacement. (**A**) Simulated surface deflection of nodes at the surface of the finger pad model for all the nine spheres. **(B)** Force-displacement relationships of the fingertip simulated for elasticity-radius combinations.

For the proprioceptive cue simulated for active touch, fingertip displacement was derived from the translational movement of the fingertip bone in the normal direction. In general, as illustrated in Fig. S3B, an increase in the radius or the elasticity will contribute to a decrease in the fingertip displacement given the same loading force.

### Geometry of the fingertip model

As illustrated in Fig. S4A, the fingertip-stimulus contact and movement was simulated with finite element models of the human distal phalange and compliant spheres. Two 2D models were built to characterize the geometry and material of the fingertip (*28*). As shown in Fig. S4B, the plane-strain model was created from a cross-sectional slice from proximal first digit to distal tip, to account for stimuli contacting across the width of the finger. As shown in Fig. S4C, the axisymmetric model was built to analyze the stimuli contact mechanics normal to the contact surface.

The finger bones and nails were modeled as rigid bodies as they were much stiffer than soft tissues. Three layers of soft tissues were included as deformable bodies, wrapping around the finger bone, namely epidermis, dermis, and hypodermis. There were no relative displacements among layer interfaces. The fingernail was modeled as 13 mm in length and 0.46 mm in thickness.

Near the surface of the finest dimension, the mesh was built with 0.25 mm wide elements. Gradually, larger sizes were used closer to the finger bone. Triangular meshes were used both in the plane-strain and axisymmetric models.

**Fig. S4.**
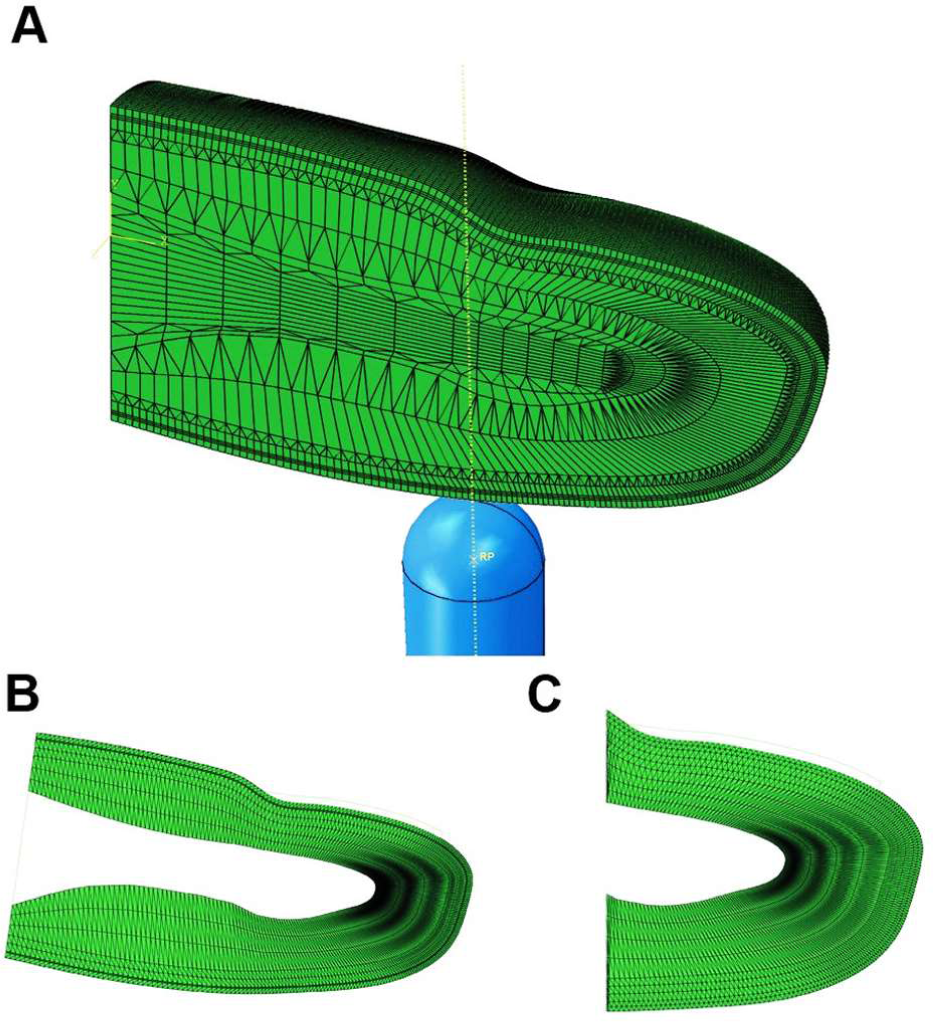
Geometry of the finger and stimulus tip model. **(A)** The compliant stimulus is implemented as hemispheres contacting the skin surface of the finger pad. (**B**) Plane-strain model to fit the surface deflection. (**C**) Axisymmetric model to fit force-displacement relation and perform simulations.

### Fitting hyperelastic material properties

In general, the plane strain model (Fig. S4B) was first utilized to fit material properties of the surface deflection, and the axisymmetric model (Fig. S4C) was then used to fit force-displacement relations and further perform simulations with compliant stimuli.

In the first step, material properties of the innermost layer (i.e., hypodermis or subcutaneous tissue) were predefined, and only the initial shear modulus of the epidermis and dermis were adjusted to calibrate the relative ratios between the elasticity of each layer. Since surface deflection is independent of the absolute material properties and is only controlled by the relative ratios between layers, deflection data from prior in vivo experiments were used to fit all the ratios in our model.

Specifically, the initial shear modulus of the hypodermis layer was set to be 1 kPa, as similar to Wu *et al*. (*42*). The reasonable search range for dermal elasticity was set as 1 to 100 times of the hypodermis, and for epidermal elasticity was 10 to 1000 times of the dermis. An exhaustive fitting procedure was applied with a step size of 0.2 on a common-log scale within this search range (Fig. S5A). Two rigid body indenters were used herein, both rigid cylindrical indenters with diameter of 3.17 and 9.52 mm, and both employing the plane strain model. Surface deflection was first simulated at six displacements (0.5, 1.0, 1.6, 2.5, 3.0, and 3.5 mm) with candidate ratios and then compared with the in vivo experimental data from Dandekar (*43*). As illustrated in Fig. S5A, the average of the coordinates of all points with a *R*^*2*^ ≥ 0.8 were then calculated as the optimal ratios. The final modulus ratio between the epidermal, dermal, and hypodermal layers was 510.63: 21.37: 1.00.

In the second step, based on fitted ratios and surface deflections, the elastic moduli of the materials were scaled to fit the displacement-force data measured for four subjects in prior experiments (*44*). Two rigid indenters were used to simulate the displacement-force responses in the axisymmetric model: a cylinder of 6.35 mm diameter, and a flat plate. By optimizing the average *R*^*2*^ for each indenter per each subject using the L-BFGS-B algorithm, the optimal scaling for each material layer was determined. As illustrated in Fig. S5B and detailed in Table S1, the final *R*^*2*^ was 0.99 and final shear moduli were 1.21 MPa for epidermis, 50.67 kPa for dermis and 2.37 kPa for hypodermis (*28*). This result is comparable to prior data by Wu *et al*. (*42*).

**Fig. S5.**
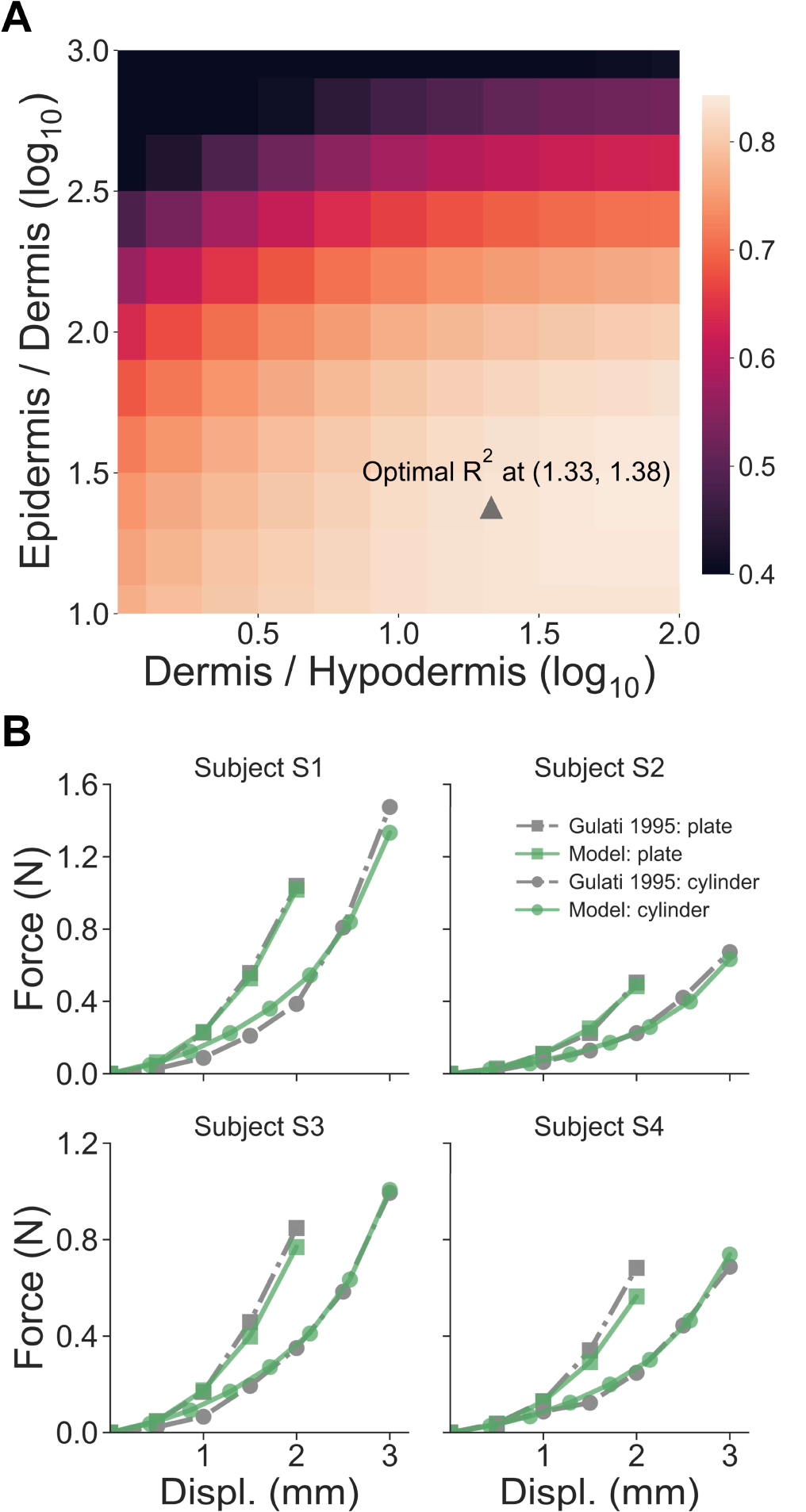
Results of the material properties fitting. **(A)** Relative ratios between skin layers are optimized to fit the surface deflection simulated by the model. The optimal point is selected by averaging all points with a R^2^ ≥ 0.8. (**B**) Force-displacement fits between model simulations and experimental measurements.

In the second step, based on fitted ratios and surface deflections, the elastic moduli of the materials were scaled to fit the displacement-force data measured for four subjects in prior experiments (*44*). Two rigid indenters were used to simulate the displacement-force responses in the axisymmetric model: a cylinder of 6.35 mm diameter, and a flat plate. By optimizing the average *R*^*2*^ for each indenter per each subject using the L-BFGS-B algorithm, the optimal scaling for each material layer was determined. As illustrated in Fig. S5B and detailed in Table S1, the final *R*^*2*^ was 0.99 and final shear moduli were 1.21 MPa for epidermis, 50.67 kPa for dermis and 2.37 kPa for hypodermis (*28*). This result is comparable to prior data by Wu *et al*. (*42*).

**Table S1.** Material properties derived from the fitting. The final shear moduli for the skin layers are taken as the average of all subjects’ results.

| Subject | Epidermis (MPa) | Dermis (kPa) | Hypodermis (kPa) | $R^2$ |
| --- | --- | --- | --- | --- |
| #1 | 1.74 | 72.74 | 3.40 | 0.99 |
| #2 | 0.83 | 34.61 | 1.62 | 0.99 |
| #3 | 1.31 | 54.96 | 2.57 | 0.99 |
| #4 | 0.97 | 40.48 | 1.90 | 0.97 |
| **Final Modulus** | 1.21 | 50.67 | 2.37 | 0.99 |

### Perceptual cues measured from one representative participant

In the biomechanical measurement experiments, gross contact areas were measured using the ink-based procedure in both passive and active touch. As illustrated in Fig. S6A-B, in both interaction conditions, the illusion case spheres (i.e., 10 kPa-4 mm, 90 kPa-6 mm, and 90 kPa-8 mm) indeed generate similar contact areas, as opposed to the distinct stimulus 10 kPa-8 mm, which generated significantly higher contact areas.

**Fig. S6.**
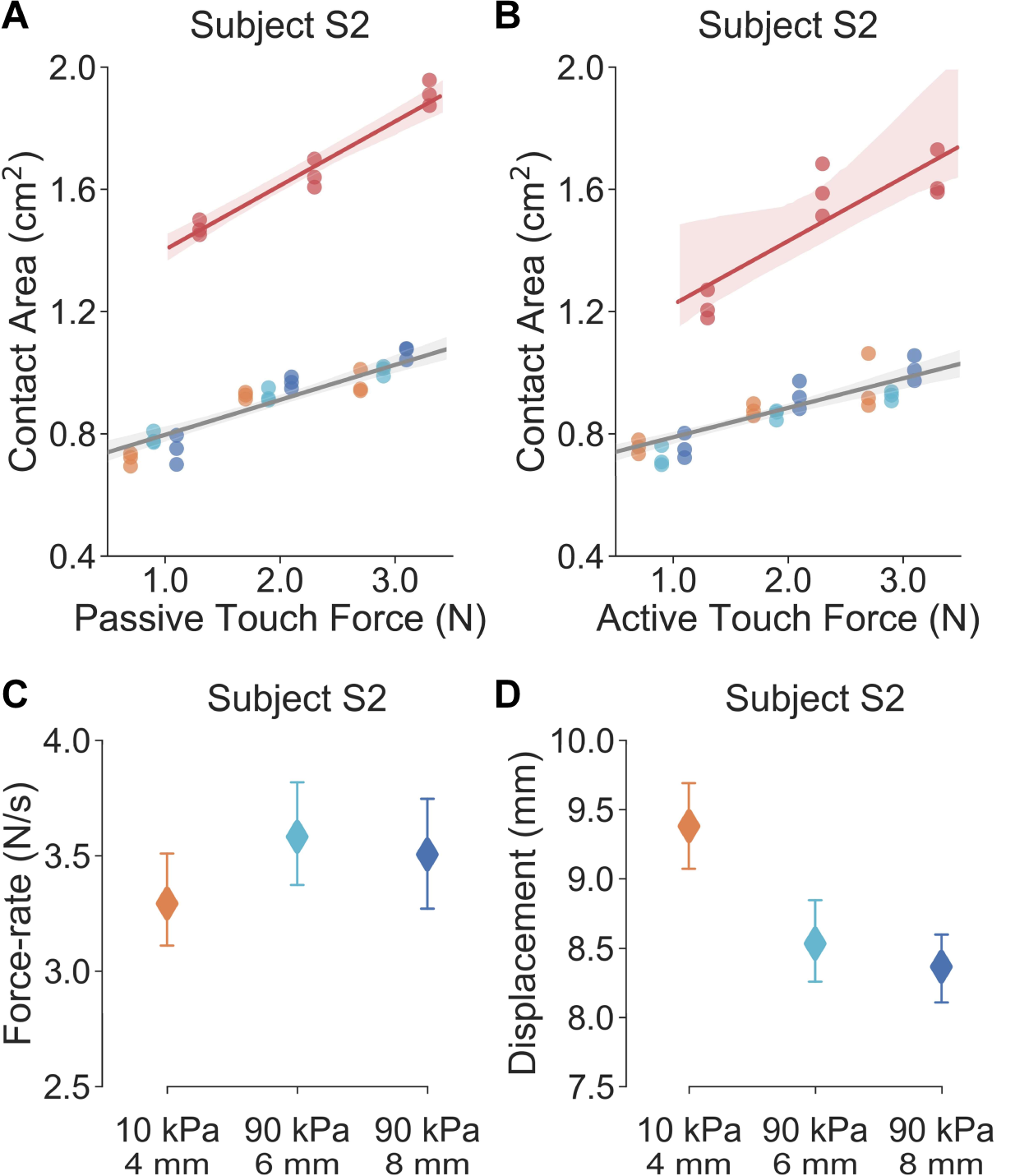
Perceptual cues measured in human-subjects experiments. Gross contact areas measured in passive (**A**) and active touch (**B**) from one representative participant. Linear regression procedures are applied to visualize the correlation between touch force and contact area. Translucent bands denote 95% confidence intervals for regression estimations. (**C**) Similar force-rates are volitionally controlled and applied in active exploration of illusion case spheres. (**D**) Distinct fingertip displacements are applied in discriminating the illusion case spheres.

Furthermore, a strong linear correlation was verified between touch force and contact area. In passive touch (Fig. S6A), the Spearman’s rank coefficient yielded linear correlations of 0.90 (*p* = 2.43*e*^-10^) and 0.95 (*p* = 9.58*e*^-5^) for the illusion and distinct spheres respectively. In active touch (Fig. S6B), the Spearman’s rank coefficient yielded linear correlations of 0.89 (*p* = 4.67*e*^-10^) and 0.84 (*p* = 0.004) for the illusion and distinct spheres respectively.

This indicates that within the designated force range, the change of contact area, elicited by the touch interaction, was proportional to the touch force. Since the force-rate was linearly controlled in passive touch, the change in contact area could be directly quantified by the force-rate cue within current measurement limitations. This results also explained that the force-rate cue induced in passive touch indeed elicited the change of contact area, which exhibited a strong linear correlation and is consistent with prior work (*5, 20*).

**Table S2.** Results of psychophysical evaluations for all stimulus pairs. Percent correct responses for each stimulus pair under different experimental conditions with all participants aggregated.

| **Experimental Conditions** | **Discriminability (correct / total)** |  |  | **Agg.** |
| --- | --- | --- | --- | --- |
|  | **10, 4 – 90, 6** | **10, 4 – 90, 8** | **90, 6 – 90, 8** |  |
| Active same force-rate | 66/90 | 77/90 | 83/90 | 83.7% |
| Passive same force-rate | 25/60 | 34/60 | 24/60 | 46.1% |
| Passive inverse force-rate | 33/60 | 35/60 | 27/60 | 52.8% |
| Passive direct force-rate | 44/60 | 45/60 | 49/60 | 76.7% |
| Summary for passive touch | 56.7% | 63.3% | 55.6% | / |
| **Aggregation** | 62.2% | 70.7% | 67.8% | / |

## Notes

### Competing Interest Statement

The authors have declared no competing interest.

